# A Phylogenomic Study Quantifies Competing Mechanisms for Pseudogenization in Prokaryotes

**DOI:** 10.1101/224238

**Authors:** Eliran Avni, Dennis Montoya, David Lopez, Robert Modlin, Matteo Pellegrini, Sagi Snir

**Author notes:** Equal contribution (Matteo Pellegrini, Sagi Snir).

## Abstract

**Background:** Pseudogenes are non-functional sequences in the genome with homologous sequences that are functional (i.e. genes). They are abundant in eukaryotes where they have been extensively investigated, while in prokaryotes they are significantly scarcer and less well studied. Here we conduct a comprehensive analysis of the evolution of pseudogenes in prokaryotes. For this analysis we consider a broad range of bacteria, but also focus on the leprosy pathogen *Mycobacterium leprae*, which contains an unusually large number of pseudogenes which comprise approximately 40% of its entire genome.

**Results:** We have developed an informatics-based approach to characterize the evolution of pseudogenes. This approach combines tools from phylogenomics, genomics, and transcriptomics. The results we obtain suggest the presence of two mechanisms for pseudogene formation: failed horizontal gene transfer events and disruption of native genes.

**Conclusions:** We conclude that while in most bacteria the former is most likely responsible for the majority of pseudogenization events, in mycobacteria, and in particular in *M. leprae* with its exceptionally high pseudogene numbers, the latter predominates. We believe that our study sheds new light on the evolution of pseudogenes in bacteria, by utilizing new methodologies that are applied to the unusually abundant *M. leprae* pseudogenes and their orthologs. As such, we anticipate that it will be of broad interest to both evolutionary biologists as well as microbiologists.

## 1 Introduction

Pseudogenes are inactive copies of genes that are functional in other contexts [40, 27]. A functional copy of a pseudogene may reside either within the same organism, suggesting it is a paralog of the pseudogene, or in other organisms, in which case they are orthologs [10]. Pseudogenes are more prevalent in eukaryotes where genome compactness is less critical and their loss confers minor fitness benefits to the organism [15]. They also serve as a reservoir for future genes and functionality. Therefore pseudogenes play an important evolutionary role.

The term pseudogene was first coined by[16] as part of their research on the *Xenopus laevis* genome. In fact, for many years pseudogene studies were mostly focused in eukaryotes where the phenomenon is prevalent[33, 38, 24]. By contrast, in prokaryotes pseudogene were generally regarded as rare [23]. Consequently, considerably less effort was devoted to pseudogene research in prokaryotes. Nevertheless, the existence of mycobacteria with large numbers of pseudogenes suggest tha pseudogenzation may also play an important evolutionary role in prokaryotes. Moreover, the formation of pseudogenes may differ between prokaryotes and eukaryotes, in part due to differing rates of horizontal transfer.

The intracellular pathogen *Mycobacterium leprae (M. leprae*), the causative of the skin disease leprosy, provides an extraordinary model to study gene loss, or reductive evolution [15]. *M. leprae*’s genome includes 1133 pseudogenes and 1614 protein-coding genes [4, 14]. The pseudogene content of *M. leprae* is significantly higher than most other bacteria, and also higher than the closely related *Mycobacterium tuberculosis*[23]. It is likely that within the intracellular environment of the host cell in which *M. leprae* grows, these pseudogenes are dispensable, and therefore the introduction of premature stop codons in these genes does not lead to a fitness loss [36]. From the study of *M. leprae* we can attempt to understand why these genes are no longer necessary for the pathogen to thrive within its host.

A phylogenomic analyses in which individual gene histories are constructed and compared across bacteria can be used to characterize the evolution of pseudogenes. While the history of speciation is normally perceived as a vertical ancestor-descendant process, which translates to a *tree-like* structure, in prokaryotes, the vertical signal is partly obscured by the massive influence of horizontal gene transfer (HGT) [7, 29]. HGT creates a widespread discordance between evolutionary histories of different genes, where each gene evolves along its own *gene tree*. Thus, the Tree of Life (TOL) concept has been questioned as an appropriate representation of the evolution of prokaryotes [13, 19]. Nevertheless, prokaryotic evolution is primarily tree-like, and we often distill the tree-like signal from all the conflicting signals.

In light of the above discussion, in this work we perform a comprehensive phylogenomic study of pseudogenes formation, over both broad and narrow phylogenetic ranges, where the focus of the narrow range is on *M. leprae*. More precisely, we contrast the origin of pseudogenes in the narrow range to the broad range. The motivation for such a study stems from the finding by Liu et al. [23] according to which a substantial portion of pseudogenes result from failed HGTs. By contrast, the prevailing assumption for pseudogene formation in *M. leprae* is the *degradation scenario* in which a native functional gene becomes non functional as a result of a disruption. For example, Gómez-Valero et al. [14] estimated that most *M. leprae* pseudogenes were created in a relatively short time interval, which led them to conclude that they were caused in a single event.

However, they also presupposed that each pseudogene originated from a functional gene. Similar results were obtained by Babu [26]. Singh and Cole [36] investigated several degradation scenarios but also state that “The mechanism by which pseudogenes arise is not yet known”, while Dagan et al. [6] studied pseudogenes as the outcome of gene death in obligate symbiotic bacterial pathogens (while HGTs were assumed to be negligible and therefore ignored). To the best of our knowledge the relative contribution of HGT or gene degradation to the evolution of pseudogenes has never been fully characterized.

Motivated by the discussion above, here we assess the relative contribution of HGT and degradation to the formation of pseudogenes across bacteria. To address this question we use novel HGT-oriented tools, and the results are confirmed by other previously developed HGT methods. We discuss the case of *M. leprae*, with its exceptionally high fraction of pseudogenes, and use functional genes as a control group to search for distinct characteristics between the two groups. Pseudogenes and functional genes are contrasted using homologs both from broad and narrow phylogenetic ranges. To generate our evolutionary inferences over narrow ranges, we compare *M. leprae* pseudogenes to their orthologs in other mycobacteria, paying particular attention to *M. tuberculosis* due to its close evolutionary relationship. We confirm and augment our phylogenomic results with insights from other evolutionary footprints such as gene synteny and functional enrichment, as well as expression assays. Thus, our work presents a comprehensive view of pseudogene formation, including the role of the two different mechanisms that are responsible for pseudogenization - the degradation process and failed HGT.

## 2 Results

We report three results regarding the evolutionary analyses of pseudogenes. In all these analyses we compared two sets of genes: genes (or rather *COGs* - clusters of orthologous genes) whose *M. leprae* ortholog is a pseudogene, and genes whose *M. leprae* ortholog is functional (it is important to note that we analyzed only genes with orthologs in *M. leprae)*. We looked for distinct characteristics between these two sets by investigating several different evolutionary markers. We note that the word “orthologs” may sometimes be misused. According to [12], and more conspicuously to a later clarification by the same author [11], a distinction is made between orthologs and xenologs. Specifically, orthologs are two homologs whose common ancestor lies in the last common ancestor of the taxa from which the two homologs were obtained, while xenologs are two homologs whose history, since their common ancestor, involves horizontal gene transfer. Since we conclude that some pseudogenes are the result of failed horizontal gene transfer, it is possible that some so-called orthologs may in fact be xenologs.

### 2.1 Phylogenetic Analysis

Our first approach focused on a phylogenetic analysis of the genes. Our goal here was to see if by using the plurality approach, as in [42], but once for the pseudogenes and once for the non-pseudogenes, we obtain a significant difference. We focused on the species that were studied in [23] and built all relevant gene trees as follows. A preliminary step was to identify, through the Mycobrowser database [17, 22], all *M. leprae* orthologs in *M. tuberculosis*. Using Mycobrowser we divided these *M. tuberculosis* orthologs into those that are annotated as functional in *M. leprae* and those that are not, henceforth non-pseudogenes (NPGs) and pseudogenes (PGs) respectively. More specifically, pseudogenes were determined as inactive reading frames with functional counterparts in *M. tuberculosis* [5] or *M. avium*[14]. For every such *M. tuberculosis* gene, we constructed its COG using all its orthologous genes in the taxa set of [23]. We identified orthologs of *M. leprae* in *M. tuberculosis* since COGs were determined using EggNOG [30], which only includes sequences of amino acids, and therefore does not contain information about *M. leprae* pseudogenes. Next we constructed the gene tree over each COG (see Methods). This resulted in two sets of gene trees, derived from PGs and NPGs, where each tree contains a gene (leaf) of *M. tuberculosis*. We decomposed the two gene trees sets into their constituent quartets and for every 4-taxa set found its plurality quartet (see Methods). The latter yielded the plurality quartet (QP) set. We also kept the underlying set of all quartets (QA).

In order to examine whether the PG and NPG sets are distinct, we constructed a maximum quartet compatibility (MQC) tree for each set. We remark that while in [42] only eleven species were analyzed, here we interrogate a significantly larger set of sixty-four species. As MQC is computationally expensive, we used the heuristic quartet MaxCut (QMC) approach [2]. Additionally, we constructed the respective 16S rRNA tree as a robust approximation to the species tree (see Methods for more details on all these procedures).

The above procedure yielded five trees over the same taxa set: the QP trees both for PGs and NPGs, the QA trees both for PGs and NPGs, and the 16S tree. We compared these trees to test whether there were significant difference between them. Tree similarity was measured using Qfit [9], that computes the percentage of identical quartets between two trees, and was found in previous studies of ours [2, 35] to be more robust than other measures for tree comparisons. The resulting scores appear in Table 1(a).

By the theoretical result of [34], the plurality quartets should be consistent with the species tree. As we do not have the species tree, the convention is to use one or more ribosomal genes, normally the 16S rRNA gene, as a proxy. We aimed to measure the similarity of each of the quartet-based trees to the species tree proxy. As can be seen from Table 1(a), the results show that PG and NPG trees are equally distant from the 16S tree. Of particular interest are the similarities between the species tree - the 16S tree - to all four other trees (bottom line in the table). We expected these numbers to show a difference between the two gene sets (pseudogenes versus non-pseudogenes), or between the different approaches (plurality quartets versus all quartets) or any combination of them. However, we could not identify any significant difference between them. None of the quartet based trees brought us significantly closer to the species tree than the other quartet based trees.

One may hypothesize that this result is due to the similarity among the quartet trees themselves, and that therefore the dissimilarity to the 16S tree is due to the methodology we use. However, these quartet-based trees also exhibit significant dissimilarity among themselves. We note however that there are several other, possibly artefactual, reasons for this dissimilarity. First, the genes cataloged as pseudogenes are indeed pseudogenes in *M. leprae*, however in all other species,they are functional. Secondly, it is possible that the level of HGT is high, reducing any signal (as might be suggested by the fact that the trees are significantly different from the 16S tree). Finally, as indicated by several other studies (e.g. [39]), even conserved genes like the 16S are subjected to HGT, obfuscating the central trend of evolution.

**Table 1:**
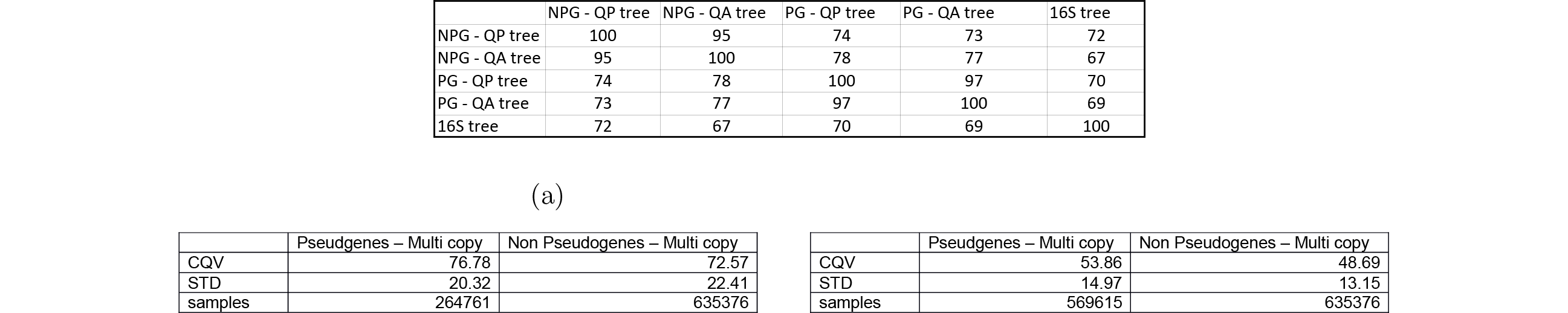
Phylogenetic Data. (a) Tree Similarities: The percentage pairwise Qfit similarity measure between the 5 trees: the QP trees both for PGs and NPs, the QA trees both for PGs and NPs, and the 16S tree. (b) CQV data: The CQV score, representing the average plurality vote over all plurality quartets. Below is the estimated STD, and number of samples (quartets). (c) CQV data: The CQV score for gene trees that may contain gene duplications, representing the average plurality vote over all plurality quartets. Below is the estimated STD, and number of samples (quartets).

|  | NPG - QP tree | NPG - QA tree | PG - QP tree | PG - QA tree | 16S tree |
| --- | --- | --- | --- | --- | --- |
| NPG - QP tree | 100 | 95 | 74 | 73 | 72 |
| NPG - QA tree | 95 | 100 | 78 | 77 | 67 |
| PG - QP tree | 74 | 78 | 100 | 97 | 70 |
| PG - QA tree | 73 | 77 | 97 | 100 | 69 |
| 16S tree | 72 | 67 | 70 | 69 | 100 |

(a)
|  | Pseudogenes – Multi copy | Non Pseudogenes – Multi copy |
| --- | --- | --- |
| CQV | 76.78 | 72.57 |
| STD | 20.32 | 22.41 |
| samples | 264761 | 635376 |

(b)
|  | Pseudogenes – Multi copy | Non Pseudogenes – Multi copy |
| --- | --- | --- |
| CQV | 53.86 | 48.69 |
| STD | 14.97 | 13.15 |
| samples | 569615 | 635376 |

Due to the lack of detectable signal in the tree distances, we resorted to another tool that is also based on quartet analysis, but may reveal other properties. For a plurality quartet topology, its *quartet vote* is the percentage of votes it obtained from all the votes for this 4-taxa set. For example, if 60% of the gene trees agree with the plurality quartet, then its quartet vote is 60%. Since we have three different topologies for a 4-taxa set, the sum over these three topologies votes is 100% and the vote of the plurality quartet is at least one-third. Now, for a set of gene trees 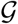 we define the *composite quartet vote* (CQV) as the average quartet vote over the entire quartet set. We can easily observe that under no HGT, every quartet in every gene tree will have a topology identical to its topology in the species tree, yielding a vote of 100% for that plurality quartet. As this holds for every quartet (4-taxa set) we will have a CQV of 100% for our gene tree set 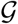. In contrast, if every gene tree is a random tree (this can be obtained by e.g. assigning a random permutation to the leaves of the gene tree), it can be easily observed that for every quartet, each of the corresponding three topologies occurs at equal probability (1/3) and so its vote will be 1/3. Thus, in this case, the CQV will be 1/3. We therefore see that the CQV ranges between 1/3 and 1 for high levels of HGT activity (where
HGT rates are infinite) and for no HGT at all, respectively. The CQV provides a summary of the intensity of HGT among the species under study (we note though that the donors of the HGTs need not be from the studied species set and even the recipients can be ancestors of the studied species, corresponding to ancient HGTs).

Therefore, seeking distinguishing characteristics between the PG and NPG gene sets, we compared their CQVs. Applying the CQV measure to the two quartet plurality sets, pseudogenes and non-pseudogenes, we obtained the scores shown in Table 1(b). The scores indicate a difference of 4.21 between the two CQVs. Normally, estimating the significance of this difference is based on an approximation that relies on the central limit theorem. However, this approximation requires that the quartet votes of different plurality quartets are independent random variables, which they are generally not. Instead, we used the Kolmogorov-Smirnov (KS) test, to check if the quartet votes, based on the pseudogenes and on the non-pseudogenes, are distributed in two statistically significant different ways. Indeed, the calculated KS statistic (0.16) was much greater than the threshold needed (0.006) to reject the null hypothesis of equal distribution with a confidence level of 99%. Combined with our previous computation showing the difference between the two CQVs, we believe this result suggests that *M. leprae* pseudogene orthologs are less prone to HGT than non-pseudogenes across the broad spectrum of bacteria we examine.

We remark that we repeated this test for a different set of gene trees, based on the same species set, and divided into pseudogenes and non-peudogenes using the same Mycobrowser classification as before, but this time allowing gene duplications. This set of gene trees is interesting since studying it may provide a more complete picture of gene evolution, while maintaining the general trend of the species phylogeny. The results of this test are presented in Table 1(c). Despite the expected decline in the CQV score (when multiple copies of a gene are represented in a tree, the number of conflicting quartet topologies is expected to increase), a similar difference between the two CQVs equal to 5.17 was found, and a KS statistic of roughly 0.16 suprpassed the 0.003 threshold indicative of a statistically significant difference between the two distributions of quartet votes. For more information, see supplementary text.

It is important to note that we searched for evidence corroborating our claim using other tools as well. Specifically, we computed the number of HGT events needed to transform the 16s tree (hypothesized to represent the correct underlying species phylogeny) to each one of the gene trees. The results of this computation, performed using RIATA-HGT [28] are presented in Figure 1. For each pair of trees, comprised of the 16s tree and one other pseudogene tree (or non-pseudogene gene tree), we plotted a blue circle (or red dot), representing the result of the RIATA computation. We then plotted the linear regression line that best suits the pseudogenes data (in magenta). Since it is clear that the putative number of HGT events is dependent on tree size, in order to determine if our collection of pseudogenes is unique in any way, we sampled 10000 subgroups from our entire collection of gene trees, while insuring those subgroups have the same distribution of tree sizes as the pseudogene trees. The linear regression lines corresponding to those subgroups are plotted as dark blue lines. Since roughly 80% of the blue lines pass below the regression line of the pseudogenes (and roughly 20% above), this test does not offer sufficient evidence to support a claim of an unusual HGT pattern in the set of psedogenes. We propose two possible explanations to this inconclusive outcome. First, the fact that RIATA is a heuristic which attempts to solve a computationally intractable problem (specifically, to minimize the number of SPR operations - Subtree Pruning and Regrafting - that are necessary to convert a species tree into a gene tree). Second, that the species tree itself may be inaccurate. This, coupled with the tree distance comparisons that were also inconclusive (Table 1), emphasizes the need to use methods that are not dependent on reference trees.

**Figure 1:**
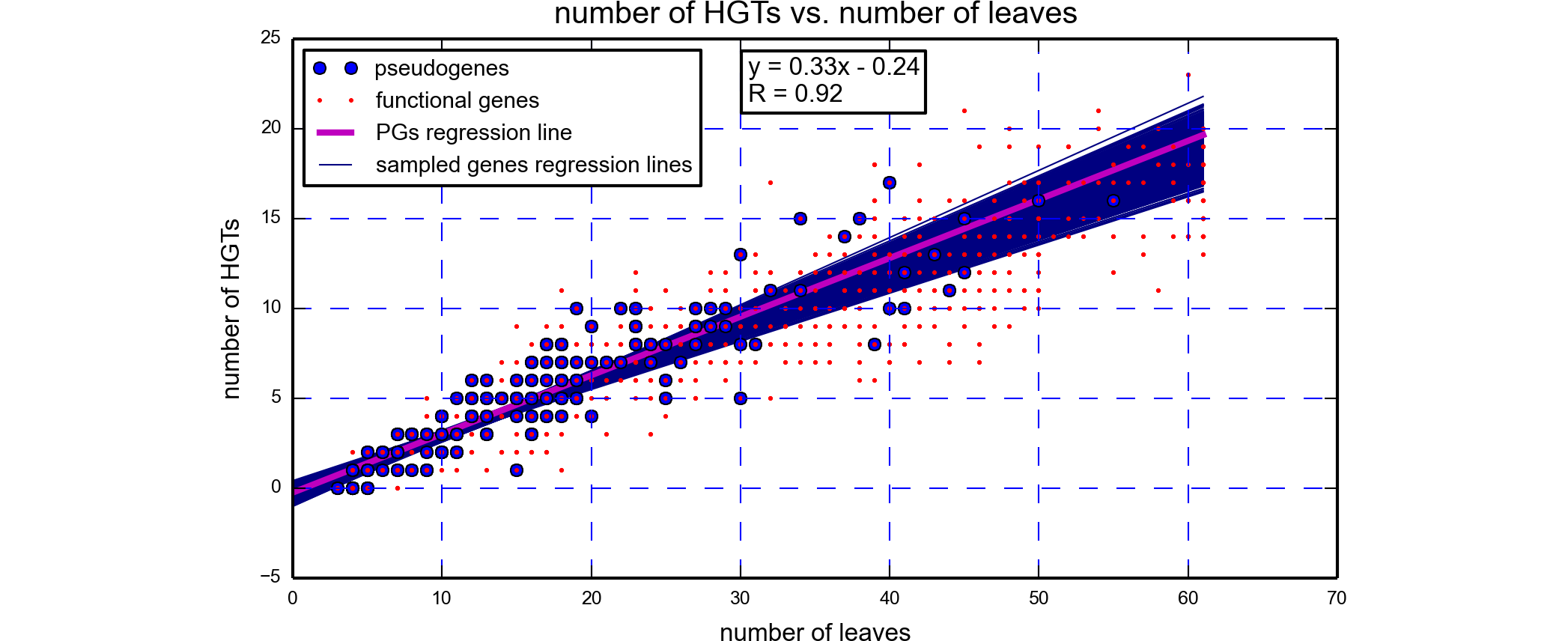
RIATA results. The parameters of the linear regression line of the pseudogenes are incorporated in the figure. Examining the number of putative HGTs vs. the number of leaves in the gene trees does not lead to an identification of an unusual HGT pattern in the pseudogenes.

### 2.2 Synteny Analysis

While the tools applied in the previous part are effective mainly for large taxa sets, and preferably with a large evolutionary span (in particular the 16S gene), in smaller and closer sets, where the signal is less pronounced, their effectiveness diminishes. We now report on another evolutionary difference between the two gene sets – pseudogenes and non-pseudogenes that we measured only between mycobacterial species. The tool we employ here to analyze the gene sets, the synteny index (*SI*, see expansion on the procedure in the supplementary text and [1, 35]) relies on a different evolutionary signal that is useful when analyzing closely related taxa.

In order to check whether the two gene types exhibit different patterns of mobility and hence also different patterns of synteny, we pursued the following procedure. To conduct an SI study between two genomes, an *orthology mapping* is necessary. In such a mapping, genomes are lists of genes, where the genes are ordered by their physical location on the genome and pairs of orthologs are connected. Briefly, a *k-neighborhood* of a gene is the set of genes at distance at most *k* from it along the genome (i.e. at most *k* genes upstream or downstream). The *k*-SI value of a gene common to two species is the portion of common genes in the two k-neighborhoods of this gene in both genomes. Clearly, the more genes those neighborhoods have in common, the greater the *k*-SI value is. In this paper we fix *k* = 10 and simply refer to the SI value.

As *M. tuberculosis* is related to *M. leprae*, most of the pseudogenes in *M. leprae* still exist in *M. tuberculosis*. Hence, we built a “star of orthology mapping” where we placed the *M. tuberculosis* genome at the center of the star and mapped four other genomes to it. In this star center *(M. tuberculosis* genome) we marked genes that are pseudogenes in *M. leprae*. Therefore, we obtained four pairs of genomes such that in all pairs, one genome is *M. tuberculosis* in which the *M. leprae* pseudogenes’ orthologs are marked (it is important to note that in *M. tuberculosis* these marked genes are non-pseudogenes). We complemented each of the genomes with genes for which no orthologs were found. We then computed the average synteny index (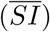) in the fashion described in Methods. However, instead of averaging over the whole genome, we averaged once over the pseudogenes and once over the non-pseudogenes, hence obtained two values of 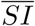 (and this is for every pair of genomes in our star). We applied this procedure to the four data sets corresponding to *M. tuberculosis* vs. (1) *M. leprae*, (2) *M. smegmatis*, (3) *M. bovis*, and (4) *M. marinum*. In order to obtain significance value for the numbers obtained we followed the same approach as in the CQV above (assuming 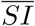 distributes normally). The results (two 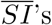) and the *p*-values (natural log values) appear in Table 2. As can be seen, the p-values obtained for all four data sets were highly significant (all smaller that *e*^-11^).

**Table 2:** **Pseudogenes vs. non-pseudogenes** 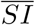 **difference:** Average synteny index results for pseudogenes and non-pseudogenes with confidence (natural log) values. Four pairs of species were compared: *M. tuberculosis* vs. (1) *M. leprae*, (2) *M. smegmatis*, (3) *M. bovis*, and (4) *M. marinum*.

| data set | #PG | SI(PG) | #n-PG | SI(n-PG) | p-val |
| --- | --- | --- | --- | --- | --- |
| M. lep | 911 | 0.63 | 1410 | 0.71 | -12.72 |
| M. smeg | 627 | 0.43 | 1226 | 0.55 | -20.39 |
| M. bov | 887 | 0.70 | 1379 | 0.78 | -11.68 |
| M. mar | 752 | 0.54 | 1308 | 0.65 | -15.38 |

The numbers in the table show a consistent trend of lower synteny for pseudogenes, that is very significant in all four pairs. Therefore, under our assumption of non-homologous recombination the neighborhood of an acquired gene in the recipient differs from its neighborhood in the donor, and we can infer that orthologs of *M. leprae* pseudogenes are more inclined to HGT than orthologs of functional genes. As detailed below, this suggests that while orthologs of *M. leprae* pseudogenes are not more inclined to HGT over a broad phylogenetic spectrum, as we discussed in the previous section, over shorter evolutionary timescales the trend is reversed.

We note that the above analysis is based on the assumption that low SI is an outcome of HGT. However, low SI can result from events of duplication as well, complicating the interpretation of our conclusions. In the supplementary text we describe the procedure we used to discard the possibility of a low SI value caused by gene duplications.

### 2.3 Functional Analysis

In this section we focus solely on *M. leprae* genes and extend our analysis by showing a relationship between expression levels of the pseudogenes and their synteny values (based on a comparison with *M. tuberculosis*). As synteny, or lack thereof, is evidence for a recombination event at a gene, this measure will lead to a grouping of pseudogenes into two classes - high SI and low SI, and we regard these as native genes versus alien genes, respectively. To examine whether there is a functional consequence to this grouping of the pseudogenes, we used expression data from human leprosy skin lesions to measure the in vivo *M. leprae* ŧranscriptome This allows a unique perspective of the expression profile of *M. leprae* in its natural host at the site of disease. Expression data was extracted as described in Methods. The results (Figure 2(a)) show the trends for both pseudogenes and non-pseudogenes. The green curve represents expression data for non-pseudogenes and serves as a control for the pseudogenes data.

**Figure 2:**
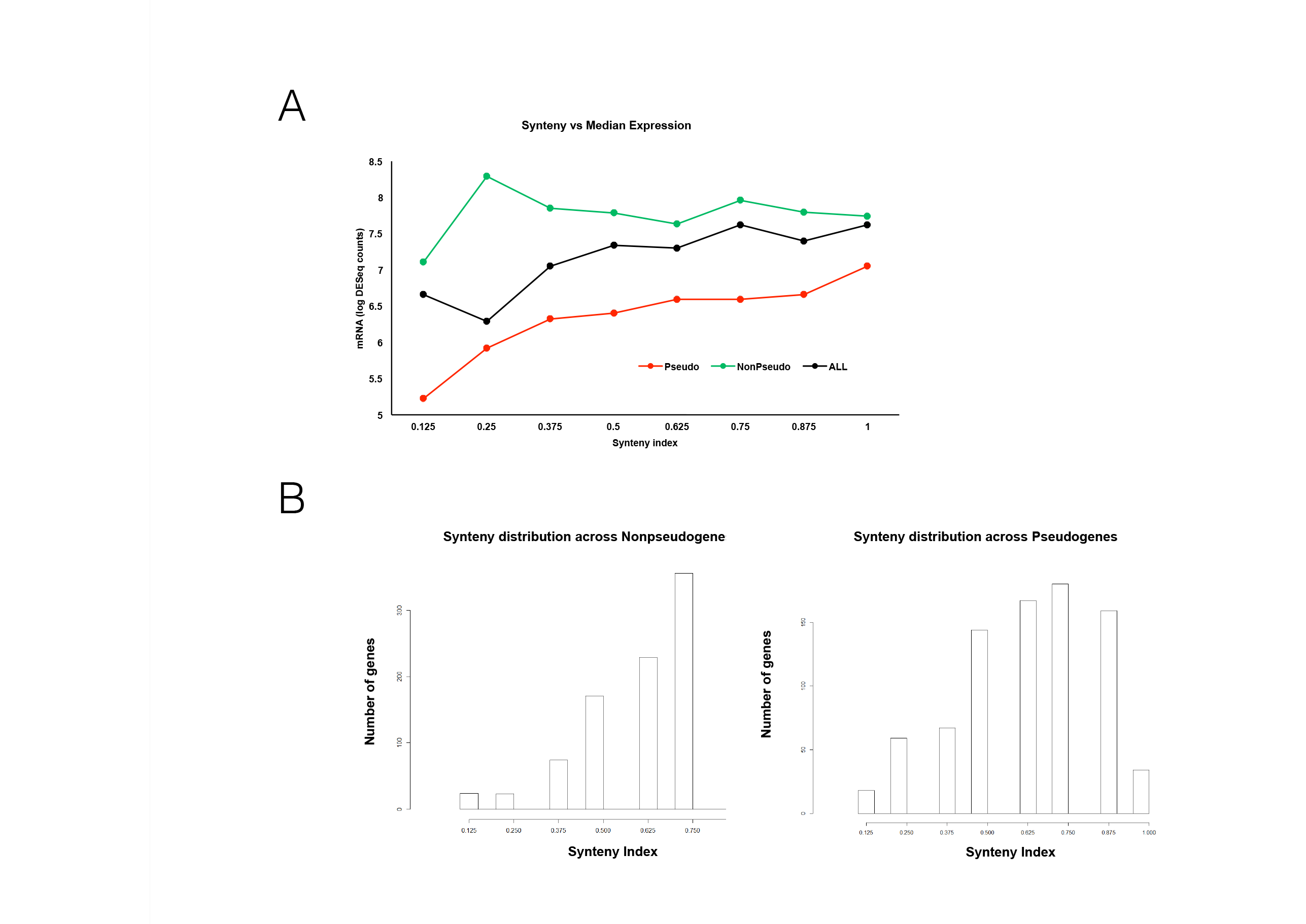
(a) **Gene Expression vs. Synteny Index**: Synteny index values between the *M. leprae* and *M. tuberculosis* genomes were calculated and compared to median gene expression in pseudogenes, non-pseudogenes, or all genes ateach synteny index value, (b) **SI Distribution at Pseudogenes and non Pseudogenes**: Gene SI distribution as measured between *M. leprae* and *M. tuberculosis* divided to pseudogenes (right) and non-pseudogenes (left).

It is evident from the figure that the expression level of functional genes is largely unaffected by their degree of synteny (except for a small decrease for very low synteny). This is expected as these genes are functional and are expressed regardless of whether they were acquired by HGT or are native in the genome. In contrast, for pseudogenes we can see a strong association between expression and synteny, and hence between expression and HGT (recall that we used SI as a proxy to HGT).

The expression profile of these low SI genes, shows that they are lowly expressed, therefore we conclude that these are failed HGT pseudogenes. However, Figure 2(b) indicates that these failed HGT pseudogenes are relatively infrequent. By contrast, the other pseudogenes, that are assumed to be native in *M. leprae* due to their higher synteny indeces, comprise the majority of pseudogenes in *M. leprae*. The latter are expressed at modest levels as they became non functional as a result of a mutation. The conclusion arising from these two results is that failed HGTs comprise the minority of pseudogenization events and are very lowly expressed, while mutation-origin (vertical evolution) pseudogenes are the majority in *M. leprae* and expressed at slightly higher levels.

We next asked how these results relate to gene functions. Previous analyses have shown that *M. leprae* pseudogenes tend to be enriched for genes involved in lipid metabolism [36]. This was consistent with the notion that the lack of *M. leprae* lipid metabolism genes caused by the formation of pseudogenes was compensated by the lipid rich environment of the host macrophage in which *M. leprae* resides. Therefore we analyzed the frequency of different functional categories in both PG and NPG, and further separated these into high and low synteny subsets. In Figure 3 (and Table 3), we show the results of this analysis for two families relating to metabolism: intermediate/respiration versus lipid metabolism. We find that PGs are enriched for lipid metabolism compared to NPGs and that this trend is even more significant for genes with high synteny. Recall that low synteny indicates HGT and conversely high synteny genes tend to be native genes. This result supports our expectation that lipid metabolism in *M. leprae* is carried out by the macrophages of the host in which the mycobacteria reside, leading to a loss of native lipid metabolism genes that become pseudogenes due to mutations. On the other hand, the same cannot be said about intermediary metabolism, where we find that PGs are depleted with respect to NPGs. In addition, further examination of Table 3 shows that cell wall and cell processes genes are more abundant in the functional genes set (in both the low SI and the high SI categories) and that regulatory proteins genes are unusually prevalent in the low SI category of functional genes. It is possible that, compared to free living bacteria, cell wall and cell processes genes in *M. leprae* are more critical to help the bacillus cope with threats from the immune system of its host. Finally, we notice that there are 76 low SI pesodugenes and only 56 low SI functional genes (Table 3). This fact suggests that the transfer of such genes (low SI) has a higher likelihood of becoming non-fuctional rather than functional, fitting with the notion that the low synteny subset corresponds to a failed HGT process [23].

**Figure 3:**
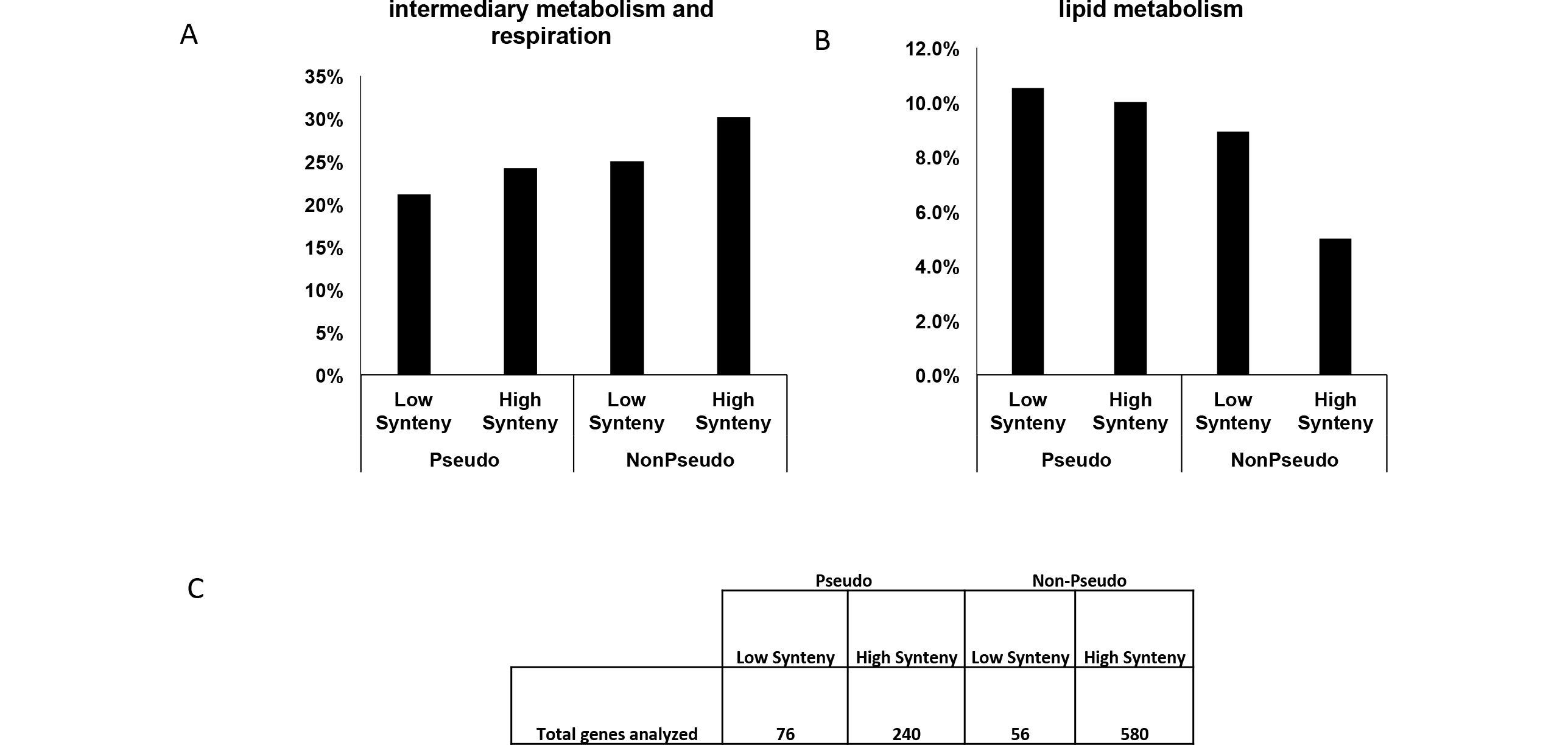
**Functional analysis of PGs and NPGs:** Percent of pseudogenes and nonpseudogenes, divided by low (0.125–0.375) and high (0.625–1.0) SI, that are part of (a) intermediary metabolism and respiration and (b) lipid metabolism, (c) total number of genes in each category.

**Table 3:** **PGs and NPGs Divided into Functional Categories:** Pseudogenes and nonpseudogenes, divided into functional categories and separated by low (0.125–0.375) and high (0.625–1) SI.

| Function | Non-Pseudogenes |  |  |  | Pseudogenes |  |  |  |
| --- | --- | --- | --- | --- | --- | --- | --- | --- |
|  | High Synteny |  | Low Synteny |  | High Synteny |  | Low Synteny |  |
|  | % of total | # genes | % of total | # genes | % of total | # genes | % of total | # genes |
| cell wall and cell processes | 25% | 147 | 32% | 18 | 19% | 45 | 24% | 18 |
| intermediary metabolism and respiration | 30% | 175 | 25% | 14 | 24% | 58 | 21% | 16 |
| conserved hypotheticals | 17% | 101 | 11% | 6 | 14% | 33 | 16% | 12 |
| N/A | 0% | 0 | 0% | 0 | 19% | 45 | 14% | 11 |
| lipid metabolism | 5% | 29 | 9% | 5 | 10% | 24 | 11% | 8 |
| regulatory proteins | 4% | 24 | 14% | 8 | 8% | 18 | 5% | 4 |
| virulence, detoxification, adaptation | 3% | 18 | 7% | 4 | 2% | 4 | 4% | 3 |
| information pathways | 14% | 79 | 2% | 1 | 4% | 10 | 3% | 2 |
| PE/PPE | 1% | 5 | 0% | 0 | 0% | 1 | 3% | 2 |
| **Total** | **100%** | **580** | **100%** | **56** | **100%** | **240** | **100%** | **76** |

## 3 Methods

### Gene Trees and Species Tree Preparation

We denote by the *species tree* the main course of speciation events yielding a given species set. In contrast, a *gene tree* represents the (evolutionary) history of a specific set of orthologous genes. Gene trees for a set of species, may differ from the respective species tree due to *reticulation events* such as horizontal gene transfer (HGT). Gene trees for our phylogenetic study were prepared as follows: First, we searched all *M. leprae* orthologs in *M. tuberculosis* using Mycobrowser [22]. Then, for each COG this search yielded, we used EggNOG [30] to find all genes in that COG that are single copy representitives of one of the 64 species in [23]. For COGs with at least four species thus found, the relevant sequences were aligned using MUSCLE [8]. Due to the large number of genes, trees were constructed using the fast tree construction FASTTREE [31]. In order to construct a species tree we used the accepted marker for this task, the 16S rRNA gene. We found 61 relevant copies of the 16S for our 64 taxa in the RDP database [3]. Sequences were extracted and aligned using MAFFT [18] using an auto alignment strategy (the one that was chosen was L-INS-i) and constructed a tree using RAxML (version 7.0.4 [37], assuming the GTR-Γ model).

### The Quartet Plurality Method

The topology of a given 4-taxa *a,b,c,d*, induced by a given gene tree, may be one of three options – *a, b*|*c, d*, or *a, c*|*b, d*, or *a, d*|*b, c*. As a result of HGT in a certain gene, the topology of the respective gene tree changes, possibly affecting the topology of the quartets. We note that tree incongruence can result also from a sequence of duplication and loss events. However, as the rates of gene loss and gain are at least an order of magnitude greater than the gene duplication rate [20, 32], and by our procedure described above of removing from the analysis species with multiple copies in a COG, we believe our main source of incongruence is HGT. As every gene is affected by different HGT events, we normally find gene trees of substantially different topologies. Consequently, the quartets induced by these gene trees may exhibit different topologies than the original, i.e., the topology induced by the species tree. In a study of eleven species of cyanobacteria using a multitude of genes [42], the topology over a given 4-taxa set that is induced by the maximum number of gene trees, was denoted as the plurality quartet or plurality topology. In [42], the set of plurality quartets over all possible 4-taxa sets was constructed and used as input to a supertree algorithm in order to overcome the frequent occurrence of HGT in these organisms. This approach was later studied from a theoretical perspective [34]. It was shown analytically that under reasonable and prevalent assumptions regarding HGT, for any 4-taxa set, its plurality quartet is the correct topology, i.e., the one induced for this 4-taxa set by the species tree. Since any tree is uniquely defined by the set of quartets it induces, this result provides us with a theoretically sound scheme of phylogenetic reconstruction that relies on the above plurality inference rule. It is important to note that even for relatively small sets of several dozens of species, the number of gene trees required to ensure that all plurality quartets will be correct with high probability is far greater than the number of genes found in any known genome. Also, as this result is theoretical, it assumes no errors in the gene trees. In reality, we do not expect all plurality quartets to be correct and hence also the plurality quartet set is inconsistent. In this case we look for the tree that satisfies the maximum number of these quartets, i.e., the MQC tree.

### Expression Analysis

Total RNA was isolated from frozen skin biopsies from lepromatous leprosy patients at time of diagnosis, before any treatment was started. Bacterial ribosomal RNA depletion was performed on total RNA by Ribozero epidemiology rRNA removal kit (Illumina). Illumina TruSeq libraries were prepared and sequenced on a HiSeq2000 and mapped to the *M. leprae* Br4923 genome using Bowtie2 [21]. Raw gene counts were normalized using DESeq2 [25] across all samples. To determine the RNA expression per gene, the DESeq normalized counts were divided by the length in kilobases of each gene.

## 4 Conclusions

In this work we investigated several evolutionary and functional properties of pseudogenes, i.e. non-functional genes, in *M. leprae* with a special emphasis on gene mobility. The analysis was performed on orthologs of *M. leprae* genes, across both broad and narrow phylogenetic ranges. It is important to note that while the *M. leprae* copy of each pseudogene is non-functional, the orthologous copies in other species are mostly functional (i.e. non-pseudogenes). These differences across species might be related to the unique environment in which *M. leprae* thrives, the intracellular environment within human skin [36].

Our analysis is divided into three parts. In the first we carry out a broad phylogenetic analysis to show that over the entire bacterial domain, non-pseudogenes are more mobile than pseudogenes. This was inferred by the higher CQV of the non-pseudogenes with respect to the pseudogenes, extracted from gene trees over the entire bacterial domain. By contrast, in the second analysis, we use synteny as a proxy for HGT to show that among the Mycobacteria, the tendency to HGT is inverted compared to the broad phylogenetic analysis, and pseudogenes exhibit more mobility.

To understand this reversal of pseudogene mobility across different evolutionary scales, we note that our functional analysis, carried out in the third part, provides a possible explanation for this observation. *M. Leprae* PGs are enriched for lipid metabolism enzymes, a function that is compensated by the lipid rich environment of the host macrophage - the environment in which *M. Leprae* thrives in leprosy lesions. However, the opposite holds when we consider orthologs of these genes across the entire bacterial domain, as most of these bacteria thrive in extracellular environments, where presumably, lipid metabolism is critical to their survival. In these extracellular environments, lipid metabolism enzymes are likely essential and therefore we expect the respective genes to be less horizontally acquired, as was previously observed and discussed in [41]. Thus the mobility of the orthologs of *M. Leprae* pseudogenes appears to depend on whether the bacteria examined are intracellular (and thus can dispense of lipid genes) or extracellular (and require lipid genes to be intact).

The last analysis extends the second by incorporating expression data of *M. leprae* genes. While PGs are not translated into functional proteins, they are often transcribed at levels similar to functional genes (i.e. NPGs). By comparing synteny (i.e. tendency to HGT) with the expression of these genes, we can distinguish between two different processes likely responsible for pseudogene formation. In the first, the gene is acquired by HGT and hence exhibits a low SI. This gene confers no advantage to the organism and remains a non functional PG. This mechanism for PG evolution corresponds to that described in [23] associated with failed HGTs as a source for pseudogenes. As such, these genes are poorly expressed in their new genome. The other mechanism corresponds to native genes that became nonfunctional as a result of a mutation in their sequence. These genes continue to be expressed, however, the transcribed RNA is not translated due to the insertion of a stop codon mutation. Figure 2(b) shows the distribution of the SI values among PG and NPG. We see that the bulk of PG and NPG reside at the higher end of the spectrum between low SI and high SI. These largely correspond to genes inherited vertically. most PGs have high synteny and high expression levels, indicating that they have not likely been gained through HGT.

Thus, in contrast to the conclusion drawn in [23], our results show that pseudogenes formed by the first mechanism (failed HGT) are in fact a small minority of the *M. leprae* pseudogene population. The explanation to this apparent contradiction, is that [23] analyzed a wide phylogenetic range, and the pseudogenes referred to there are entirely a different gene population than the pseudogenes in our study, the orthologs of *M. leprae* pseudogenes. Thus we conclude that *M. leprae* pseudogenes, which are far more abundant than those found in any other species, likely arose from the mechanism of disruption of native genes.

